# Patterns of fish utilisation in a tropical Indo-Pacific mangrove-coral seascape, New Caledonia

**DOI:** 10.1101/455931

**Authors:** A Dubuc, N. Waltham, R. Baker, C. Marchand, M. Sheaves

## Abstract

Mangrove forests are important habitats for fish. However, their utilisation by fish, and the specific values they confer, are still not fully understood. This study describes how fish use mangrove forests in an Indo-Pacific mangrove-coral reef seascape. Sampling was conducted using underwater video cameras (UVCs) to describe spatial and temporal variations in fish assemblages across a small-scale (~ 2.5 km^2^) system, and over the tidal and lunar cycle. UVCs were deployed in the two main component habitats of mangrove forests: at the mangrove forest edge, and inside the forest (5 m from the forest edge), to establish patterns of utilisation of fish across the tidal and lunar cycle. Proximity to coral reefs had a strong influence on the mangrove fish community, as most fish recorded were reef-associated. Juveniles of 12 reef species were observed, including two species classified as vulnerable on the IUCN list, and one endemic species. Fish assemblages on the mangrove edge differed significantly from those inside the forest. Most fish utilised the forest edge, with few species making regular use of in-forest habitats, supporting the contention that most fish species remain on the edge and potentially retreat into the forest for opportunistic feeding, or when threatened by larger predators. Species-specific patterns of utilisation varied across the tidal and lunar cycle. Small differences in depth profiles and substrate across the small-scale system had a significant effect on fish assemblages, highlighting the importance of accounting for spatial heterogeneity in these factors. These data provide important information for managers to implement adequate conservation strategies that include broader interconnected habitat mosaics.

## Introduction

Mangrove systems are part of a mosaic of productive coastal habitats [1] that provide a variety of services to fish and human populations [2, 3]. Mangrove forests are a fundamental component habitat of mangrove systems [4], and confer many of the attributes that make them highly valuable fish habitats [5–9]. However, studies have shown varying degree of mangrove forest utilisation, with for instance a higher contribution of reef fish species to mangrove fish assemblages in the Caribbean compared to several places in the Indo-Pacific [10–21]. These observations suggest that not all mangrove forests provide equivalent values to fish. Moreover, recent work in mesotidal Australia suggests that few fish penetrate beyond the forest boundary [15], suggesting that the use of mangrove forests is spatially heterogeneous. This new evidence raises the question relating to the specific ways in which mangrove forests are utilised by fish. More studies are needed to characterise fish assemblages in mangrove forests with different settings (coastal, estuarine, island, embayment), different tidal ranges (micro-, meso- or macrotidal), proximity of other high value habitats such as seagrass beds and coral reefs, or climatic zones [21–24]. A better understanding of how mangrove forest utilisation varies spatially and temporally would provide new insights to help explaining the contrasting results found in the literature.

In many parts of the Indo-Pacific, the tidal range is greater than in the Caribbean, where mangrove forests are usually continually available to fish [3]. Intertidal mangrove forests are challenging environments, most notably because they are only available to most aquatic organisms while they are flooded at high tide [1, 24, 25]. The intermittent availability of mangrove forests may explain the low use by fish in the Indo-Pacific [23]. Indeed, tidal variation (extent, duration and frequency of flooding) generates a range of constraints for fish utilising mangrove forests. Most evident is the decrease in water depth and eventual drainage of the forest as the tide ebbs, forcing fish out of intertidal mangrove forest zones. Several studies have indeed demonstrated that fish undertake regular migrations in tidally driven mangrove systems, with different patterns of mangrove use according to fish species, lunar cycle (neap vs spring tide) and tidal phase (flooding vs ebbing) [15, 26–29]. Migration of fish in response to tidal movements results in substantial connectivity between the three major tropical coastal habitats: coral reefs, seagrass beds and mangrove forests [25, 30], giving rise to the idea that mangrove forests are part of a wider interconnected habitat mosaic [1]. Therefore, investigating tidal and spatial variations in fish assemblages in mangrove forests is a crucial step towards fully appreciating the value and functioning of the whole tropical coastal ecosystem.

The difficulty of sampling these habitats goes a long way towards explaining the paucity of information available on fish assemblages inside mangrove forests [20, 31]. The use of conventional techniques such as underwater visual censuses or netting techniques is restricted across much of the Australasian region where saltwater crocodiles (*Crocodylus porosus*) are common, and where dense mangrove forests reduce the efficiency of most net-based approaches [15]. Recently, underwater video has been successfully applied to study in-forest fish assemblages [15, 32, 33], most notably because it overcomes a lot of sampling issues, substantially reduces field labour intensity, and allows for high-temporal and -spatial resolution data collection simultaneously in different habitats, such as the edge and the inside of a mangrove forest [34].

In this study we deployed underwater cameras on the edge and inside a mangrove forest [15, 26] coupled with high frequency depth loggers to record spatio-temporal variations in fish assemblages in a microtidal Indo-Pacific mangrove-coral reef seascape. We identified fish species that use the mangrove forest, and used an array of exploratory data analyses and modelling techniques to describe how fish utilisation changes between the forest edge and in-forest habitats, and how fish assemblages vary across the tidal cycle.

## Materials and methods

### Study site

Our study focused on a relatively pristine mangrove forest in Bouraké, South Province of New Caledonia (21° 56.971S, 165° 59.481E; Fig 1). New Caledonia is an archipelago located in the South West Pacific, 1,500 km east of Australia. New Caledonia has the largest lagoon in the world, partly registered on the UNESCO World Heritage list. New Caledonia experiences a semi-arid to tropical climate with annual total rainfall of 1,000 mm, and a mixed semi-diurnal microtidal regime (maximum 1.8 m tidal range). Bouraké receives little freshwater inflow with no defined drainage.

**Fig 1.**
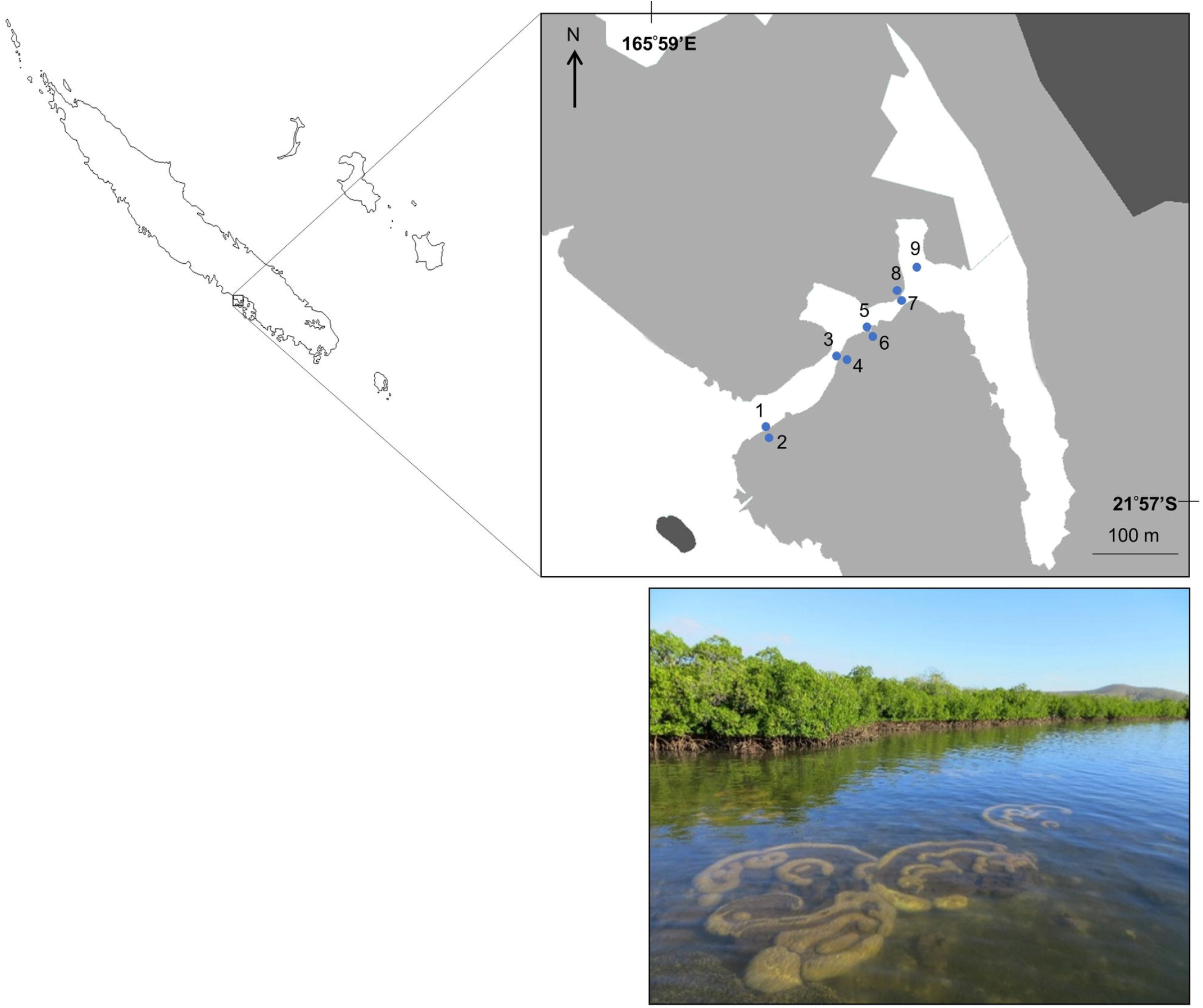
Map and picture of the study system in Bouraké, South Province, New Caledonia. The nine study sites in the mangrove channel sampled from the 21 February to 1 March 2017 are represented by their respective number. Light grey areas represent mangrove forest, dark grey areas represent mainland, and white areas represent water.

The area comprises approximately 2.5 km^2^ of mangrove forest dominated by *Rhizophora stylosa* on the seaward edge and *Avicennia marina* on the landward margin, with a large semi-enclosed central lagoon (1.2 km long, 60 m wide, 1-2 m depth). A channel (20-70 m wide, 2-6 m depth, 700 m long) connects the main lagoon to the coastal waters of Pritzbuer Bay (~ 20 km^2^). The channel comprises two sheltered inlets (approximately 0.01 km^2^ each), and a shallow (1-2 m depth) coral reef platform that extends from the middle of the channel to the edge of the mangrove forest. Corals could be seen right on the edge of the forest in some places. We chose a study location where coral reefs occur in close proximity to mangrove forests (Fig 1), effectively a seascape comparable to the Caribbean coastline [35], so we could relate our findings to this other ecoregion [36] where the tidal range is smaller.

### Data collection

Fish using the mangrove forest were examined on an inland/offshore gradient along the channel (Fig 1). To assess differences in fish assemblage composition between edge and inside the forest, 4 paired sampling were conducted (sites 1 to 8). Each paired sampling consisted of two sites within 5-7 m distance; the even site number of the paired sampling was located on the mangrove forest edge (defined as the boundary between mangrove prop-roots and bare substrate), and the odd site number located about 5 m inside the forest. Site 9 (considered an edge site in the analyses) was located on the reef platform of the innermost bay, at the edge of scattered mangrove trees slightly away from the main forest. The substrate at sites 1 and 9 consisted of dead corals, small live coral boulders and sand, while on other edge sites it comprised mainly dead corals and small and larger live coral boulders. The substrate was homogeneous and consisted of silt material at in-forest sites.

Fish assemblages were sampled using underwater video cameras (UVCs; Model ATC9K Oregon Scientific) to investigate tidal variations in fish assemblages simultaneously on the edge and inside the forest. Unbaited UVCs mounted on stable bases were deployed at each site during neap (21 to 23 February 2017) and spring tides (28 February to 1 March 2017). A sampling day consisted of cameras first deployed at sites early in the morning (first light), continuing until the battery was discharged, and, with a replacement battery, again deployed mid-afternoon at all sites until the battery was discharged (recording lasted between 2h and 2.5h). Four sampling days were completed (two during neap tides and two during spring tides). Cameras were positioned around 7 cm above the substrate, facing towards the channel. A marker mounted on a flexible rod (3 mm diameter, 0.5 m long) was placed 0.5 m in front of the camera lens as a visibility indicator to ensure a minimum visibility of 0.5 m was achieved in all videos. Visibility was very good and consistent during the sampling period, and fish could be identified confidently up to approximately 2 m from the UVCs in all videos. As depth is one of the main limiting factors to mangrove accessibility, tidal variations in water depth (cm) were measured every 15 minutes at each site with depth loggers (In-Situ Inc. Rugged Troll 100 model). James Cook University issued a permit for a limited impact research to deploy underwater cameras in New Caledonia (no endangered or protected species were involved as no collection of any specimen was conducted). The study area does not benefit from any special protection, therefore, access and activities are not restricted, and no specific permit was required to sample.

### Data extraction from videos

While UVCs allow large amounts of data to be gathered quickly in the field, considerable time is required to process these videos. Therefore, we subsampled the acquired video footage. From the two neap tide sampling days, one day was randomly selected and videos at all sites were processed for that day. For the remaining sampling day, all videos were processed from five sites; being the reef platform (site 9) and two pairs of in-forest and forest edge sites (sites 5-8). These sites were selected so one replicate for a site located on the reef platform, and two replicates of paired sites not located on the reef-platform were available. Considering this selection, the sites were then randomly chosen. The same selection was applied to the two sampling days conducted during spring tides.

Once sediments had settled after camera deployment (typically 2-3 min), videos were viewed using VLC media player (VideoLAN, 2001) and subdivided in 5-min intervals to follow tidal variations in fish assemblages. The occurrence of each fish taxon in each 5-min interval was recorded. Only presence/absence data were recorded to avoid biases induced by count data when using UVCs [15]. Fish were identified to the lowest possible taxonomic level. Features useful in discriminating species within some genera or families such as *Plectorhinchus* spp., Mugilidae spp., or Gobiidae spp. were difficult to distinguish in videos, therefore these taxa were identified to genus or family level only. When possible, juvenile fish were identified based on colour patterns and body shape. Any activity such as feeding, hiding, cruising or escaping was also noted. All fish identifications were validated by two additional experts. For each 5-min interval video processed, the information concerning the date of sampling, site, time of day, habitat (edge vs in-forest), lunar phase (neap vs spring), tide direction (flooding vs ebbing), and corresponding water depth was recorded (Appendix S1).

### Data analyses

An index depending on observation per unit effort, similar to the catch per unit effort index (CPUE) when dealing with fishing techniques, was developed to calculate frequencies of occurrence of taxa from the video data. The frequency of occurrence of each taxon was calculated per site (the total number of 5-min intervals in which a taxon was observed at a site was divided by the total number of 5-min intervals recorded at this site). Only taxa with a frequency of occurrence ≥ 0.05 at one or more sites were retained for analyses (referred to as “common taxa”). Taxa with a frequency of occurrence < 0.05 (referred to as “rare taxa”) were excluded from analyses.

Non-metric multidimensional scaling (nMDS) was used as an exploratory analysis to assess differences in fish assemblages among sites during spring and neap tides. The frequency of occurrence of each common taxon was calculated per site per lunar phase. Data were square root transformed (SQRT) to decrease the impact of extreme values, and an nMDS analysis based on Bray-Curtis dissimilarities, the most appropriate distance measure when using abundance data [37], was conducted. Clusters within the nMDS were determined by conducting an overlay cluster analysis at 40% and 45% similarity on the dissimilarity matrix of all frequencies of occurrence. A two-way analysis of similarity (ANOSIM) was used to test whether there were significant differences in fish assemblages between sites and lunar phase. Pearson correlations exceeding R > 0.7 between the ordination and taxa were used to fit vectors on the nMDS plot. All analyses were performed using PRIMER 6 software [38]. Additionally, frequencies of occurrence of each common taxon at in-forest and edge sites were calculated and plotted using horizontal bar plots to further investigate differences in fish assemblage composition between the two habitats.

To investigate the factors impacting fish presence/absence, a General Linear Mixed Model (GLMM) was conducted using the package “glmm” in R [39]. The GLMM was conducted on all the 5-min intervals recorded (Appendix S1) with presence/absence of any common taxa for each 5-min interval (1 if any common taxa were observed in the 5-min interval, or 0 if no common taxa were observed) as the response variable, “Depth”, “Habitat” (edge vs in-forest), “Lunar phase” (neap vs spring), and “Time of day” (morning vs afternoon) as the fixed factors, and “Site”, “Date”, “Tide direction” (flooding vs ebbing) and a nested effect of “Site” within “Habitat” as the random factors, using a Bernoulli distribution and a logit link function.

Cumulative depth frequency curves were plotted for each site to highlight differences in temporal dynamics. To further understand how fish utilisation varies across depth, variations of SQRT frequencies of occurrence across depth, over flooding and ebbing tide on the edge and in-forest were assessed using a General Additive Mixed Model (GAMM). Each 5-min interval was allocated to a class of water depth of 10 cm (from 10-20 cm to 120-130 cm) according to the water depth value recorded, and the SQRT frequencies of occurrence of each common taxon was calculated per class of water depth during flooding and ebbing tide per habitat (the total number of 5-min intervals in which a taxon was observed at a class of depth during flooding and ebbing tide per habitat was divided by the total number of 5-min intervals recorded for this same sample unit). Frequencies of occurrence were SQRT to reduce the impact of extreme values. To avoid false absence recordings, taxa never recorded in the habitat of interest were not considered (i.e. if a taxon was never recorded in-forest during the study it was not included in the in-forest analysis). To run the GAMM, SQRT frequencies of occurrence were used as the response variable, “Depth” as a smooth term, and “Habitat” and “Tide direction” as parametric terms using a Gaussian distribution and an identity link function. “Habitat” was included in the model to avoid any nesting issue. The model was built using the package “mgcv” in R [40]. Patterns of variations of SQRT frequencies of occurrence were then investigated graphically using boxplots to examine the variations of average SQRT frequencies of occurrence among taxa at each depth interval, and a LOESS curve was fitted to the data to analyse the general pattern of habitat use. Patterns of mangrove habitat use for each taxon were then plotted using a LOESS curve and individually assessed graphically to examine similarities and classify patterns of fish occurrence across depth. Taxa were grouped in similar patterns if their maximum average occurrence was observed at a similar tidal stage. Three equivalent tidal stages were defined for this purpose: Low tide (between 10-20 and 40-50 cm); Mid tide (between 50-60 and 80-90 cm); High tide (between 90-100 and 120-130 cm).

## Results

### Fish composition

Fifty-six video deployments were analysed (totalling more than 118h of video). Seventy-two taxa from 29 families were recorded, with 36 common taxa (frequency of occurrence ≥ 0.05 on at least one site) retained for further statistical analyses (Table 1). Most species recorded were marine and reef-associated [41] including fish of families Scaridae, Chaetodontidae, Pomacanthidae, Siganidae, Acanthuridae, Lutjanidae, or Labridae.

**Table 1.**
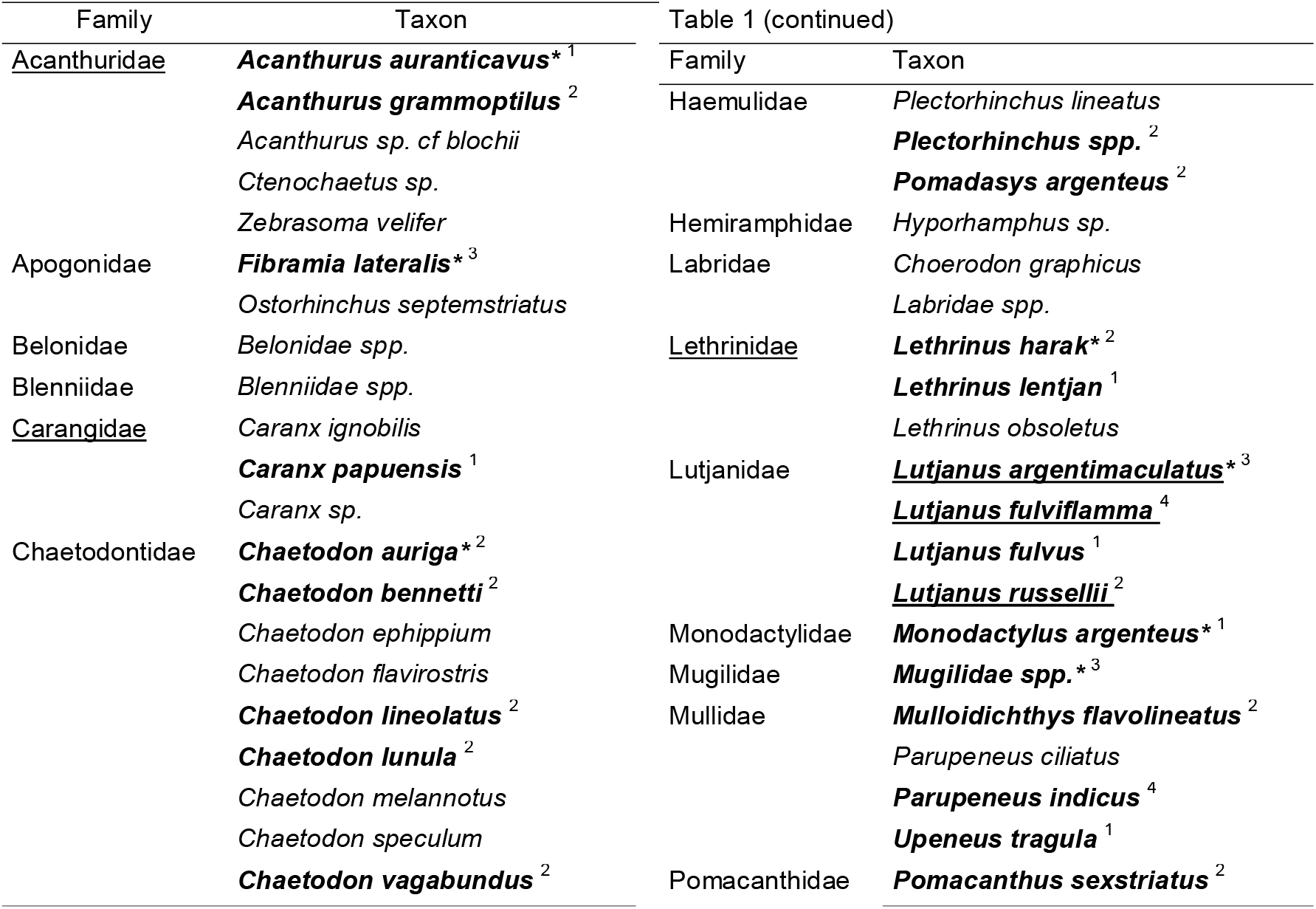
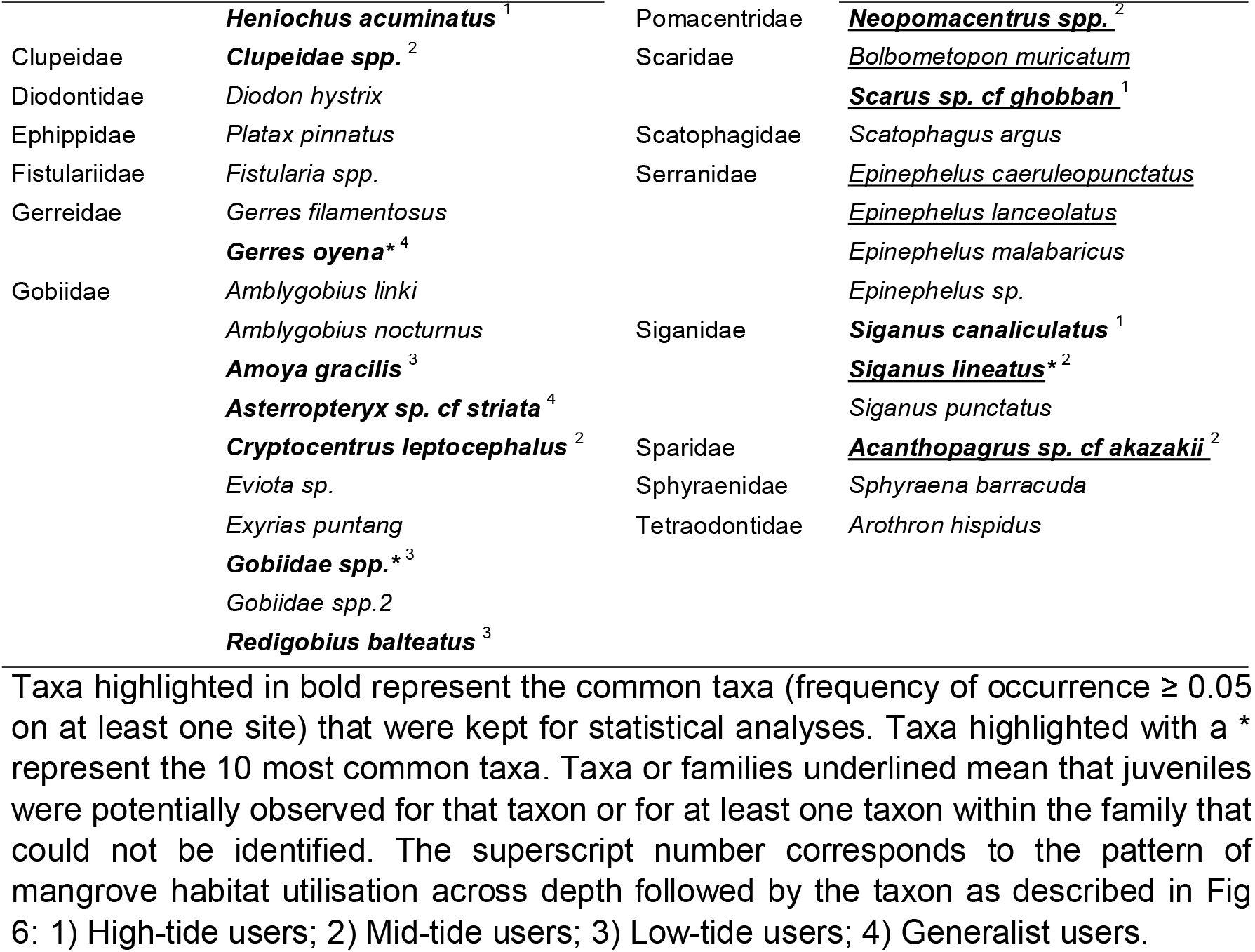
Summary of all the families and taxa identified at Bouraké, New Caledonia.

Fish composition varied significantly among sites (ANOSIM: R = 0.793, p < 0.001), with distinct assemblages generating three and four different clusters at 40 % and 45 % similarity respectively on the nMDS plot (Fig 2). At 40 % similarity, the 1^st^ cluster comprised all the samples conducted in-forest. The samples were characterised by a lower taxonomic richness (23 common taxa; Fig 3) dominated by *Fibramia lateralis* and all the taxa belonging to the Gobiidae family (except *Cryptocentrus leptocephalus* and *Asterropteryx* spp.), that were the only taxa recorded almost exclusively at in-forest sites (Fig 3). The 2^nd^ cluster comprised all the samples conducted on the edge but site 7 at spring tide. The samples were characterised by a higher taxonomic richness (34 common taxa; Fig 3), among which 10 taxa, mostly reef-associated, significantly contributed to the fish assemblage composition at edge sites (Fig 2). Site 7 at spring tide was an outlier and made the 3^rd^ cluster driven by the abnormally high occurrence of *Neopomacentrus* spp. (Fig 2). Interestingly at 45 % similarity, another cluster was generated, separating deep edge and shallow edge sites (Fig 4, Fig 2). Three species of snappers, *Lutjanus fulviflamma, Lutjanus argentimaculatus and Lutjanus russellii* were the only three species not showing apparent preference for edge or in-forest sites as they were almost evenly recorded on the two habitats (Fig 3), and therefore did not significantly characterised any of the two habitats (Fig 2). Replicate samples plotted close to each other and were grouped in the same clusters (Fig 2). Lunar phase did not significantly influence fish assemblages (ANOSIM: R = 0.2, p > 0.2).

**Fig 2.**
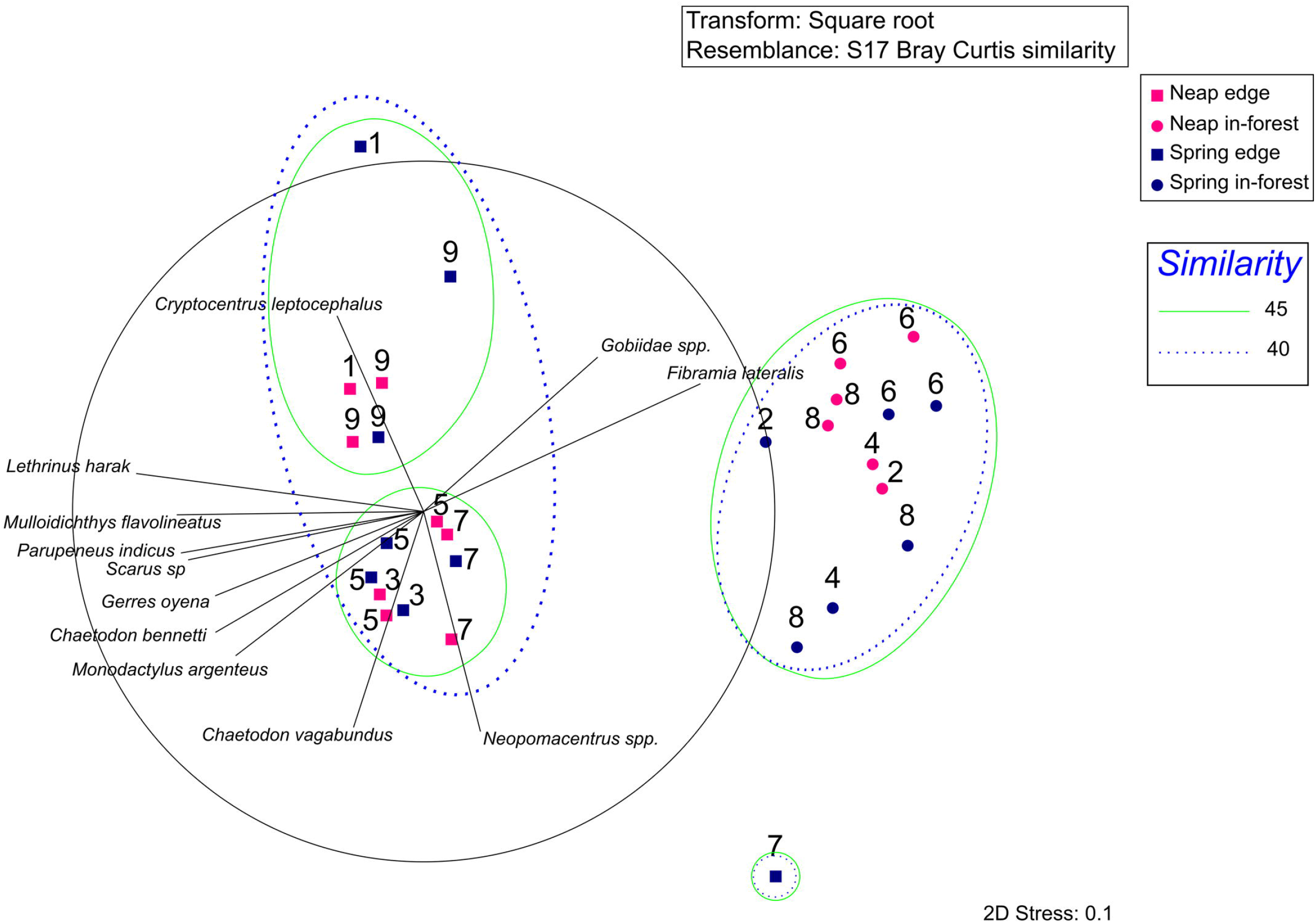
nMDS analysis performed on square root transformed frequencies of occurrence for each taxon per site per tide. Edge sites are represented by squares and in-forest sites by circles. Sites sampled at neap tide are coloured in deep pink, and sites sampled at spring tide in navy blue. Solid green and dotted blue ellipses represent overlay clusters determined at 45 and 40 % similarity respectively. Vectors represent taxa with a Pearson correlation with the ordination R > 0.7.

**Fig 3.**
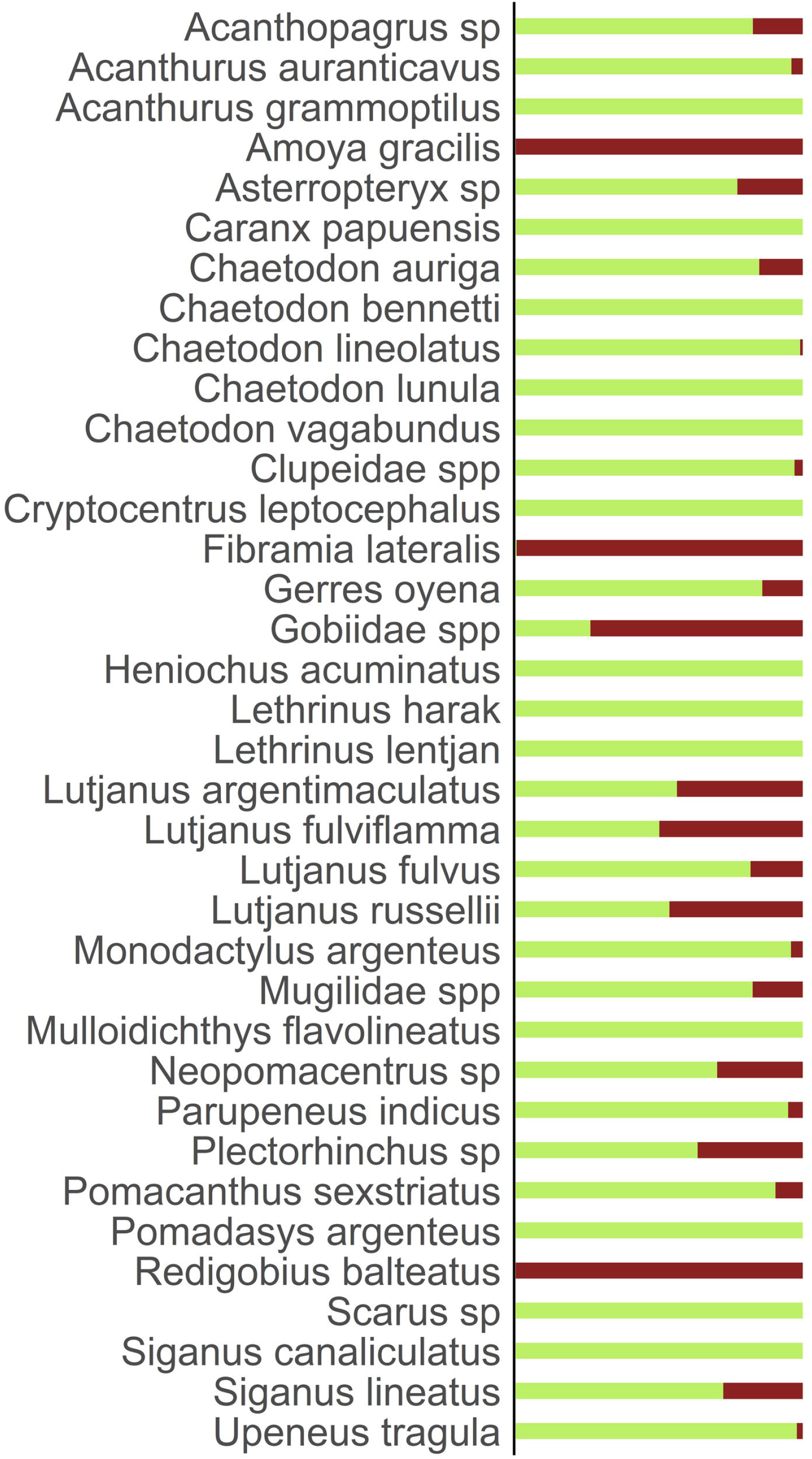
Proportion of time spent by each of the common taxa on the edge versus inside the forest. edge=green and in-forest=brown. Proportions range from 0 to 1, 1 corresponding to a taxon exclusively recorded on the edge or in-forest and 0.5 corresponding to a taxon recorded on the edge as frequently as in-forest.

**Fig 4.**
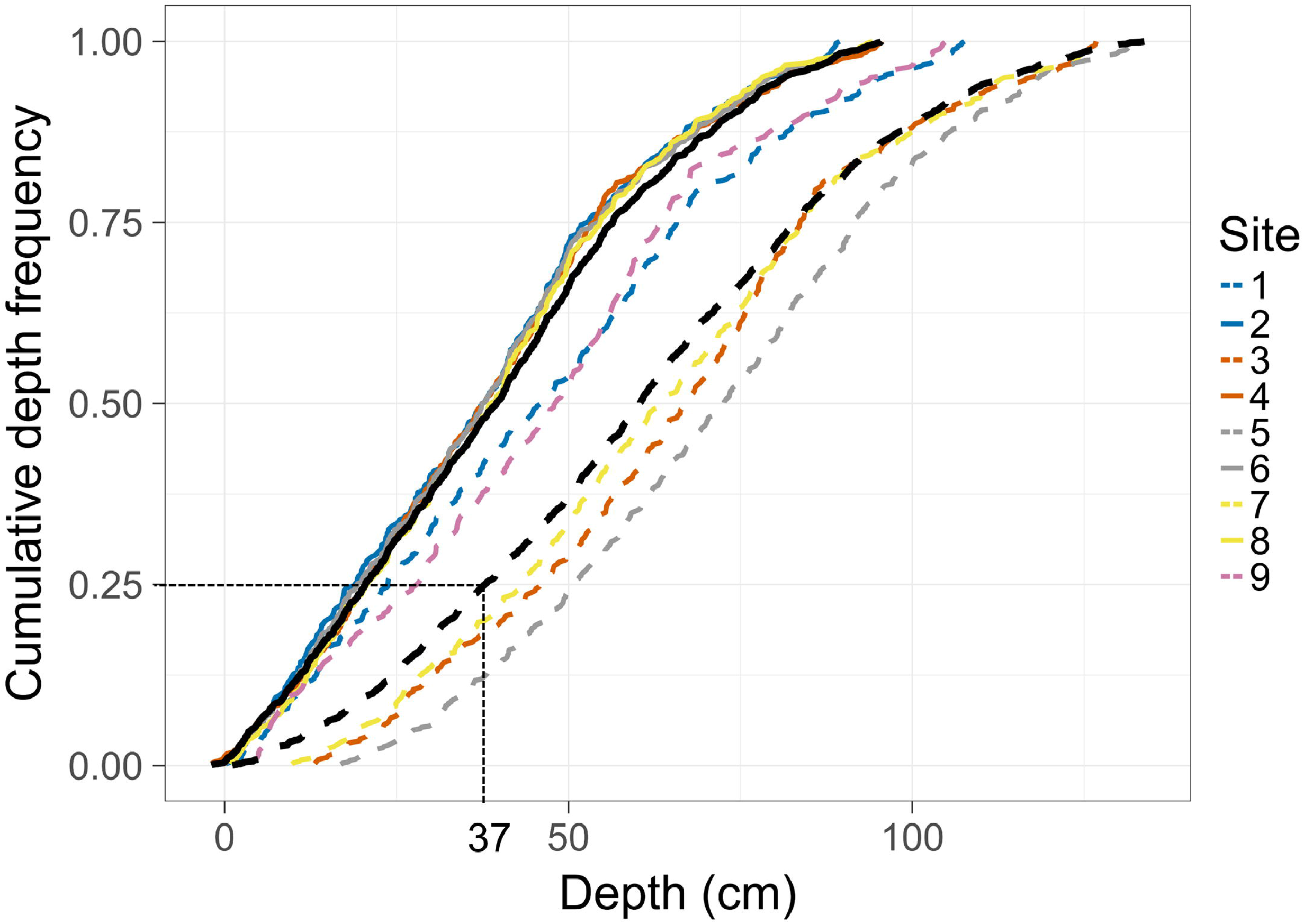
Site-specific cumulative depth frequencies. Each colour represents a paired edge and in-forest site, and edge sites are represented by dashed lines and in-forest sites by solid lines. The dashed dark line indicates the mean cumulative depth frequencies across all edge sites. The solid black line indicates the mean cumulative depth frequencies across all in-forest sites. An example is provided to help interpret the figure (for 25 % of the recorded time depth was on average equal or below 37 cm at edge sites).

“Habitat” (GLMM: z = −2.637; p < 0.005), “Lunar phase” (GLMM: z = −9.406; p < 0.001) and “Depth” (GLMM: z = −2.118; p < 0.05) significantly influenced the presence/absence of taxa. However, “Time of day” (GLMM: z = −1.519; p > 0.1), date of sampling (GLMM: z = 1.555; p > 0.05), “Tide direction” (GLMM: z = 0.991; p > 0.1) and sites within a same habitat (GLMM: z = 1.394; p > 0.05) did not significantly influence presence/absence of taxa. Further data exploration following the GLMM results showed that at edge sites there was a higher proportion of 5-min intervals in which a taxon was observed compare to in-forest sites. Similarly, during neap tides, there was a higher proportion of 5-min intervals in which a taxon was observed compare to spring tides (Appendix S2).

### Tidal variations in fish assemblages

Average depth was substantially shallower at in-forest than edge sites (neap tides (mean ± SE): 34 ± 0.57 and 55 ± 0.66 cm respectively; spring tides: 48 ± 1.11 and 71 ± 1.23 cm respectively), as was maximum depth (95 cm and 133 cm respectively; Fig 4). Moreover, in-forest sites were exposed (i.e. not flooded) for 4-5 h per day during neap tides, and 2-3 h per day during spring tides, while sites on the edge were always submerged. Sites could be classified into three groups according to depth profiles (Fig 4): deep edge sites (sites 3, 5, 7; maximum depth 133 cm); shallow edge sites (sites 1 and 9; maximum depth: 107 cm); in-forest sites (sites 2, 4, 6, 8; maximum depth: 95 cm; Fig 4).

As the GLMM showed that depth had a significant effect on presence/absence of taxa, a GAMM was used to further explore the response of frequencies of occurrence of fish across increasing and decreasing depth (equivalent to flooding and ebbing tide) and by habitat. SQRT frequencies of occurrence of fish significantly varied across depth (GAMM: F = 6.756; p < 0.001; Appendix S3), and significantly differed between “Habitat” (GAMM: F = 39.792; p < 0.001) and “Tide direction” (GAMM: F = 9.056; p < 0.005). Magnitude of variations in SQRT frequencies of occurrence across depth was higher at edge than in-forest sites (Figs 5a and 5b, Appendix S3). However, the patterns were similar between the two habitats, with overall frequencies of occurrence highest at mid-tide, especially mid-ebb tide, and lowest at extreme depth values (low or high tide; Figs 5a and 5b, Appendix S3).

**Fig 5.**
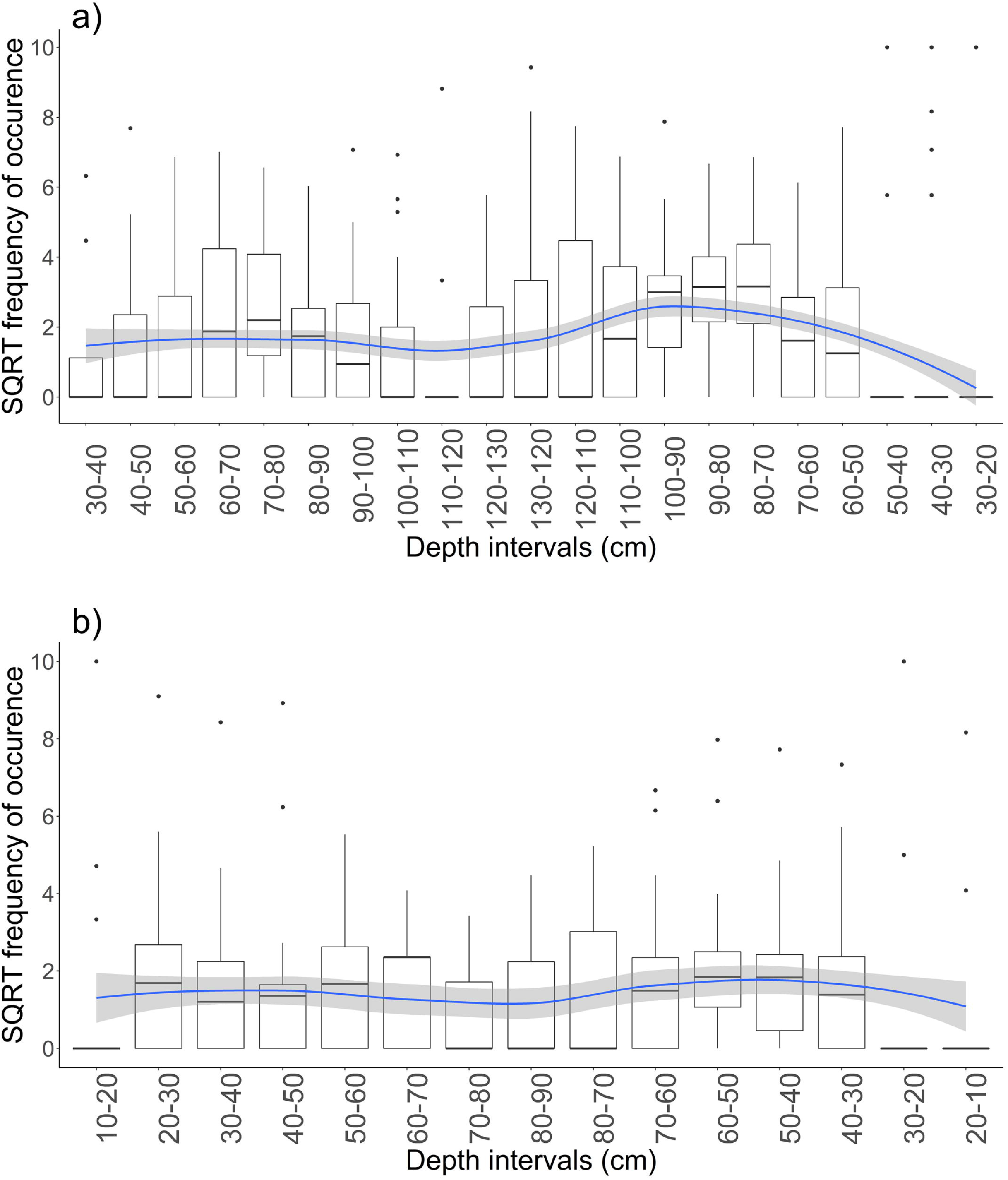
Boxplots of average square root transformed frequency of occurrence of common taxa across depth on a) edge sites; b) in-forest sites. The blue line is the LOESS curve representing the general pattern of habitat use for all common taxa considered. Shaded area around the LOESS curve represents the 95% confidence interval. On Fig 5b, interval 90-80 cm has been removed as no data were recorded.

Similarities in mangrove forest utilisation among common taxa clearly determined 4 main patterns of utilisation: 1) taxa with higher frequencies of occurrence at high tide (90-130 cm; High-tide users); 2) taxa with higher frequencies of occurrence at mid-tide (50-90 cm; Mid-tide users); 3) taxa with higher frequencies at low tide (10-50 cm; Low-tide users); 4) taxa with similar frequencies of occurrence across depth (Generalist users); Fig 6, Appendix S4).

**Fig 6.**
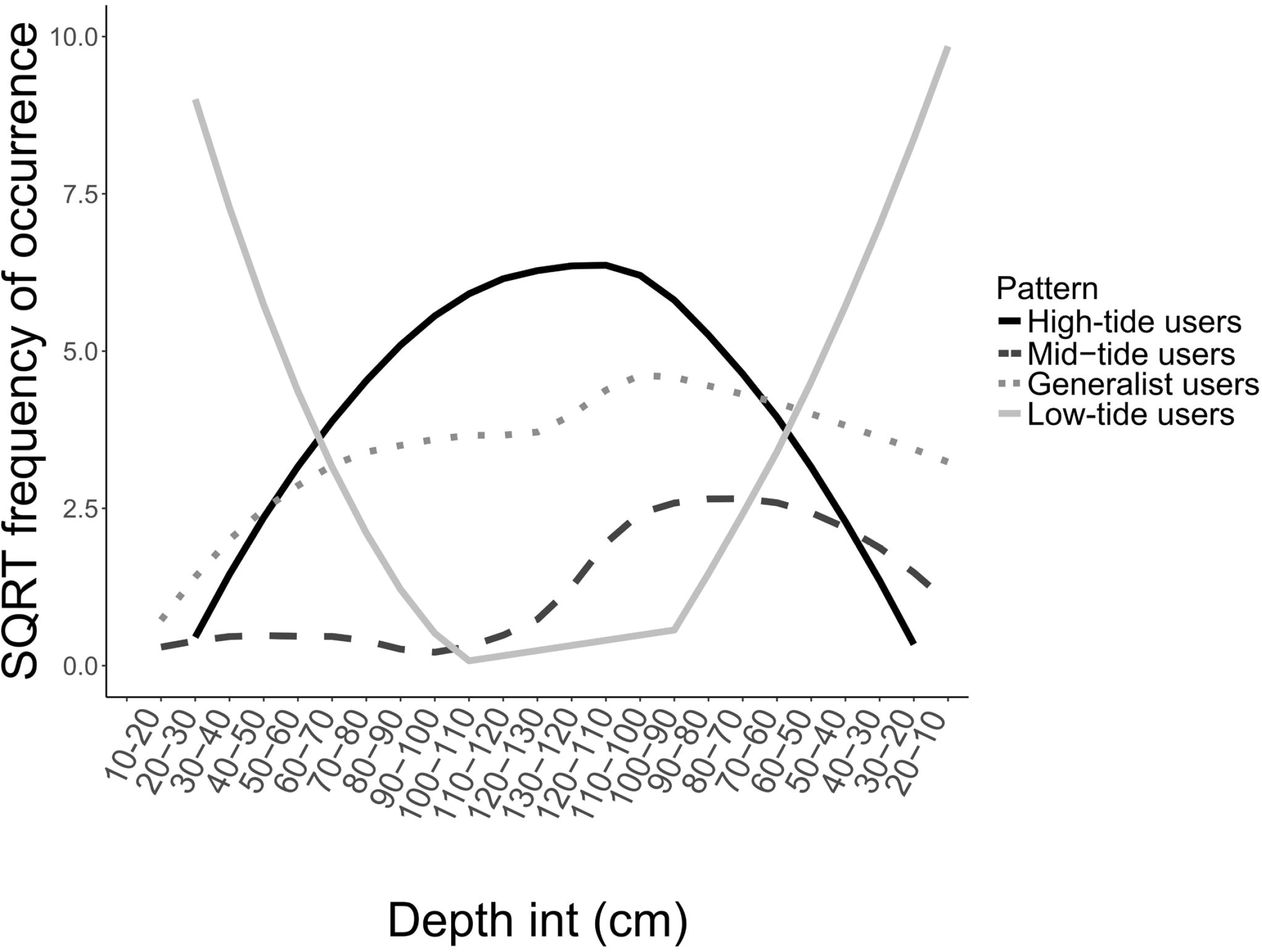
Patterns of mangrove habitat utilisation across the depth profile. The curves represent the LOESS curves constructed with the square root transformed frequencies of occurrence of fish across depth. Common taxa followed four main patterns of mangrove habitat utilisation across depth: 1) taxa using mangrove habitats at high tide (High-tide users); 2) taxa using mangrove habitats at mid-tide (Mid-tide users); 3) taxa accessing and leaving mangrove habitats at low tide (Low-tide users); 4) taxa without any apparent preferences for depth (Generalist users). Table 1 identifies the taxa allocated to each category.

## Discussion

Understanding the spatial and temporal variations in the use of mangrove habitats by fish is important when considering conservation and resource management to protect mangrove ecosystems from human and natural disturbances. This study highlights that the edge and inside of mangrove forests, the two major component habitats of mangrove forests, featured distinct taxonomic diversity and fish assemblage composition. Fish assemblages varied significantly across the tidal cycle, with species-specific patterns of mangrove habitat utilisation. Spatial differences in water depth among sites within a same habitat also seemed to influence fish assemblages across this mangrove/coral system. While only a small portion of the species observed on nearby coral reefs were recorded in Bourake, we found that this mangrove forest does have a role in supporting reef fish species, emphasising the importance of Indo-Pacific mangroves as valuable fish habitats.

The setting of this tropical mangrove/coral system influenced the nature of the fish assemblages recorded. At least 72 taxa made use of this relatively small mangrove/coral system, with most species classified as tropical marine and reef-associated [41]. Most taxa recorded have not been identified as mangrove-associated in previous studies in coastal mangroves in the west Pacific [15, 16, 42], suggesting that their presence is linked to the proximity of coral reefs, further supporting the contention that mangrove-coral habitats are interconnected. Conversely, many fish families important in other Indo-Pacific mangrove systems remote from coral reefs, such as Leiognatidae, Ambassidae, Sillaginidae, Terapontidae, or Toxotidae [15, 16, 28, 43] were not recorded in Bouraké. Most individuals observed were at a sub-adult stage, however, juveniles where occasionally recorded for several taxa. Juveniles of at least 12 reef fish species were commonly recorded (*Lutjanus fulviflamma, Lutjanus argentimaculatus, Lutjanus russellii, Lethrinus* spp. (2 species), *Bolbometopon muriculatum, Siganus lineatus, Caranx* sp., *Epinephelus caeruleopunctatus, Scarus* sp., *Acanthurus* sp., *Neopomacentrus* sp.). Additionally, relatively small individuals of *Epinephelus lanceolatus* and *Acanthopagrus akazakii* were observed. Thus, the fish community using this system consisted of a substantial number of juvenile reef species, including juveniles of two species classified as vulnerable on the IUCN list (*E. lanceolatus* and *B. muricatum*), and one endemic species (*A. akazakii*) [44]. These findings highlight that near-coral mangrove habitats in the Central Indo-Pacific, such as Bouraké, have a role in providing habitats for juvenile reef fish in parallel to the situation in the Tropical Atlantic [12, 45, 46].

While early studies concluded that high connectivity between coral reefs and mangroves had little influence on mangrove fish assemblages [18, 28, 47], recent evidence suggest that in many instances there is a high occurrence of reef-associated fish in mangroves adjacent to reefs [10, 19, 20, 48]. While supporting this idea, the current study emphasises that the utilisation and value of mangrove forests vary locally and cannot be generalised from one system to another [10, 15, 21].

This study highlighted clear spatial variations in fish assemblages across the two different habitats mangrove edge and mangrove in-forest. Indeed, fish assemblages were distinctly different between the mangrove edge and just a few meters inside the mangrove forest. Most fish were recorded cruising on the edge of the mangrove forest, while sightings inside the mangrove forest were sparser. Two main hypotheses, namely increased food supply and providing shelter, have been suggested to explain why fish use mangrove forests. However, neither of these two hypotheses were confirmed by the current study as few foraging activities were recorded and few individuals were observed actively sheltering among mangrove prop-roots. In fact, few species made regular use of the mangrove forest, supporting the idea that most fish species simply remain on the edge and potentially retreat into the forest for opportunistic feeding, or to escape presence of larger predators [7]. This result aligns with the observations in estuarine mangrove forests of northern

Australia where few species made regular use of the in-forest habitat [15]. These two habitats (edge and in-forest) probably confer different values to fish, however, fish could benefit from most attributes that physically attract them in mangrove systems [8] by using the mangrove fringe without venturing into the forest. This result supports the idea that high tidal range leading to forest drainage limits the use of mangrove forests in the Indo-Pacific compare to the Caribbean. Accessing the forest could be disadvantageous because of increased risk of becoming trapped after the tide falls, but could also be linked to adverse water quality such as low dissolved oxygen that develops at low tide [49].

Fish assemblages exhibited small-scale spatial (dozens of meters) heterogeneity, particularly along the forest edge compared to in-forest sites. There was a clear distinction in terms of fish assemblages in the nMDS plot between sites 1 and 9, and sites 3, 5 and 7. This pattern could be explained by water depth profile and substrate differences, with sites 1 and 9 featuring dead corals, small live coral boulders and sand, and shallow depth, while other edge sites also had dead corals, along with small and larger live coral boulders but lacked sand, and experienced deeper depth. Conversely, all the in-forest sites were quite similar in terms of fish assemblages, suggesting that they provide a homogeneous habitat with similar substrate and depth profile throughout the system. <u>Johnston and Sheaves (2007)</u> [50] also identified species-specific responses to different small-scale habitats according to their depth and substrate composition. The importance of accounting for spatial heterogeneity of fish assemblages when characterising the habitat value of a system, or when using fish assemblages as a bio-indicator of ecological change or ecosystem health [51], was highlighted by <u>Becker et al. (2012) [52] who</u> observed the influence of small spatial scale changes in water depth and substrate composition on fish assemblages at seagrass beds in South Africa.

Fish assemblages varied temporally across the tidal cycle. Tide-induced depth variations have been linked to changes in fish assemblages [26, 28, 52, 53]. This result was corroborated here as fish assemblages varied across depth, with more fish observed during mid-tides, especially at mid- ebbing tides, and most species generally avoiding extreme shallow or deep water. In fact, fish displayed species-specific responses to depth with four main patterns identified: 1) taxa using mangrove habitats at high tide (High-tide users); 2) taxa using mangrove habitats at mid-tide (Mid-tide users); 3) taxa accessing and leaving mangrove habitats at low tide (Low-tide users); 4) taxa without any apparent preferences for depth (Generalist users). Patterns 3 and 4 mainly comprised taxa that frequently used mangrove habitats such as *Fibramia lateralis, Lutjanus argentimaculatus, Siganus lineatus, Gerres oyena*, or taxa belonging to the Gobiidae family [15], while the other two profiles comprised mainly marine and reef-associated species. In essence, rather than accessing mangrove habitats as soon as they become available, many species seem to use mangrove habitats only for a restricted period of time. Other studies that looked at variations in fish assemblages across the tidal cycle also reported species-specific responses to the depth profile and highlighted that species using mangrove habitats extensively were accessing them at a shallower depth than other less frequently observed species [26, 52–56]. Factors driving these tidal migrations are not fully understood, and the fact that species do not enter mangrove habitats as soon as they become available may suggest that these patterns could be the result of behavioural adaptations to avoid adverse water conditions such as low dissolved oxygen that can occur early or late in the tide [49]. Species using extensively mangrove habitats could be adapted to tolerate lower depth and adverse dissolved oxygen conditions compare to species that would occasionally use mangrove habitats when they are more suitable. More studies are needed to link tidal fish migrations with dissolved oxygen conditions in mangrove habitats because dissolved oxygen is likely to be a critical environmental factor determining the value of these habitats.

Lunar phase was another influential factor responsible for temporal variations in mangrove habitats utilisation by fish. More fish were detected during neap tide than spring tide, however, taxonomic richness and fish assemblage composition were similar. These data oppose previous studies that observed more fish at spring tide than neap tide [57–59]. These authors suggest that spring tides result in more habitats available and for longer duration, attracting more fish. We firstly thought this was an artefact of the methodology, with fish disappearing from the field of view as water became too deep. However, we compared fish occurrence within the same depth intervals between neap and spring tides, and fish presence was still substantially lower during spring tides, which suggests that there may be another explanation. One explanation could be that at spring tides fish can access more intertidal habitats, reducing the probability of encounter with the UVCs. We also observed very strong currents in the channel and along the mangrove edge during spring tides that could reduce the time fish can benefit from using mangrove habitats as the energy needed to remain on the mangrove edge may be too high.

## Conclusion

The results here provide further support that within a mangrove forest, the inside and edge of the forest are two distinct habitats characterised by different fish assemblages. The study mangrove forest plays a role in maintaining a substantial number of fish species. However, the habitats use was species-specific, suggesting that utilisation and value need to be considered species-by-species if we want to fully understand the role mangrove systems play in maintaining fish communities. The high spatial and temporal heterogeneity of fish assemblages complicates the characterisation of mangrove forests value and utilisation, suggesting that results from one location are unlikely to be applicable to other systems more broadly. This is an important conclusion for managers when considering to adapt conservation strategies from other locations, to local-specific habitat mosaics.

## Acknowledgements

We thank Dr. Adrien Jacotot, Dr. Thanh Nho Nguyen and Mrs Clara Hass for their assistance in the field. We greatly appreciated suggestions from members of the Science for Integrated Coastal Ecosystem Management group to improve the manuscript.

## Supporting information

S1 Appendix. Presence/absence of all common taxa for each 5-minute interval recorded during the study.

S2 Appendix. Percentage of 5-minutes intervals with no common taxa observed at a) neap tide vs spring tide and b) edge vs in-forest habitats.

S3 Appendix. General Additive Mixed Model for a) edge sites and b) in-forest sites.

S4 Appendix. Similarities in species-specific patterns of mangrove utilisation across depth: a) High-tide users; b) Mid-tide users; c) Mid-tide users (continue) d) Low-tide users; e) Generalist users.

