## Supplementary figures and images for "Patterns of fish utilisation in a tropical Indo-Pacific mangrove-coral seascape, New Caledonia"

### Appendix S2

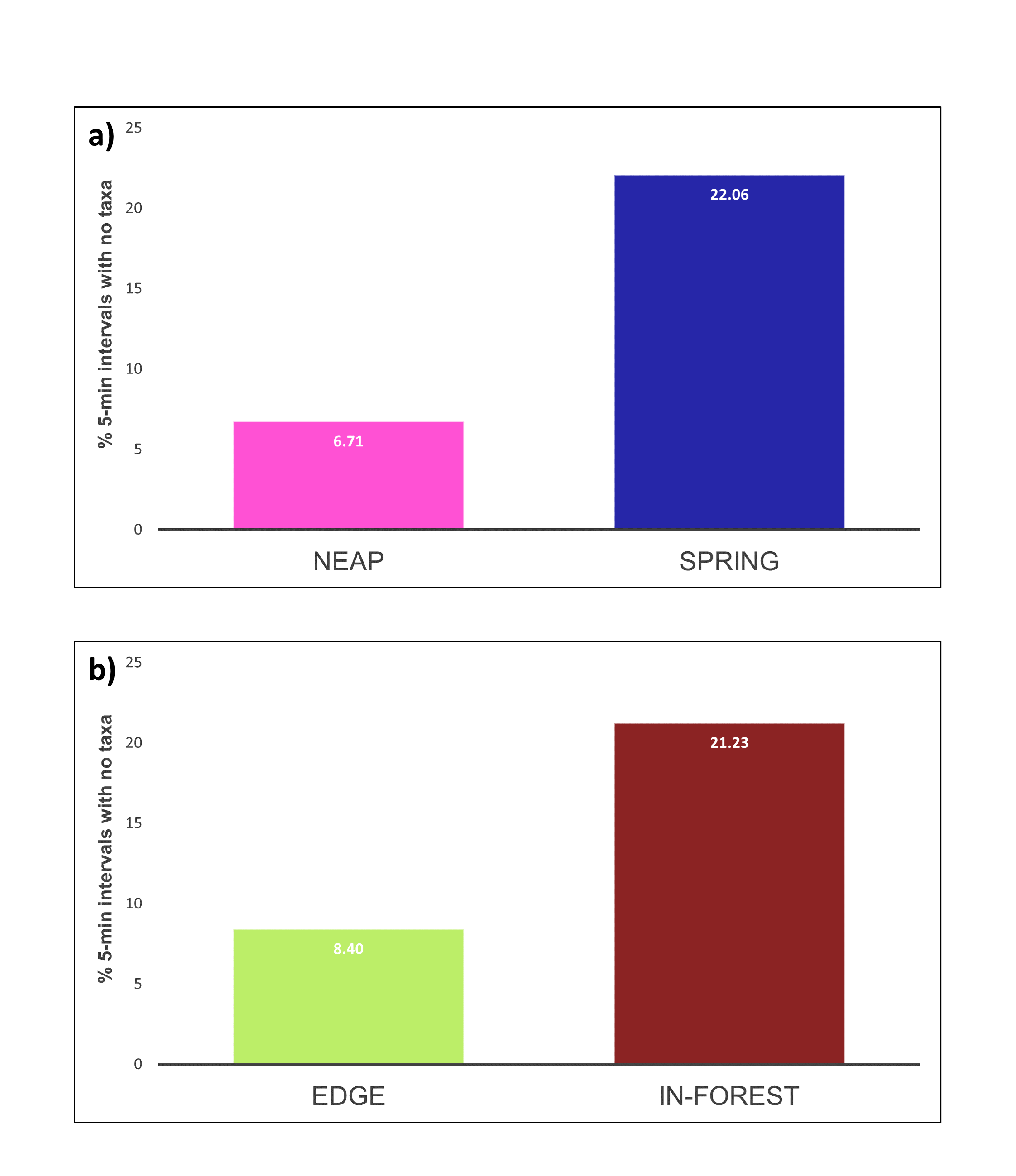

### Appendix S3

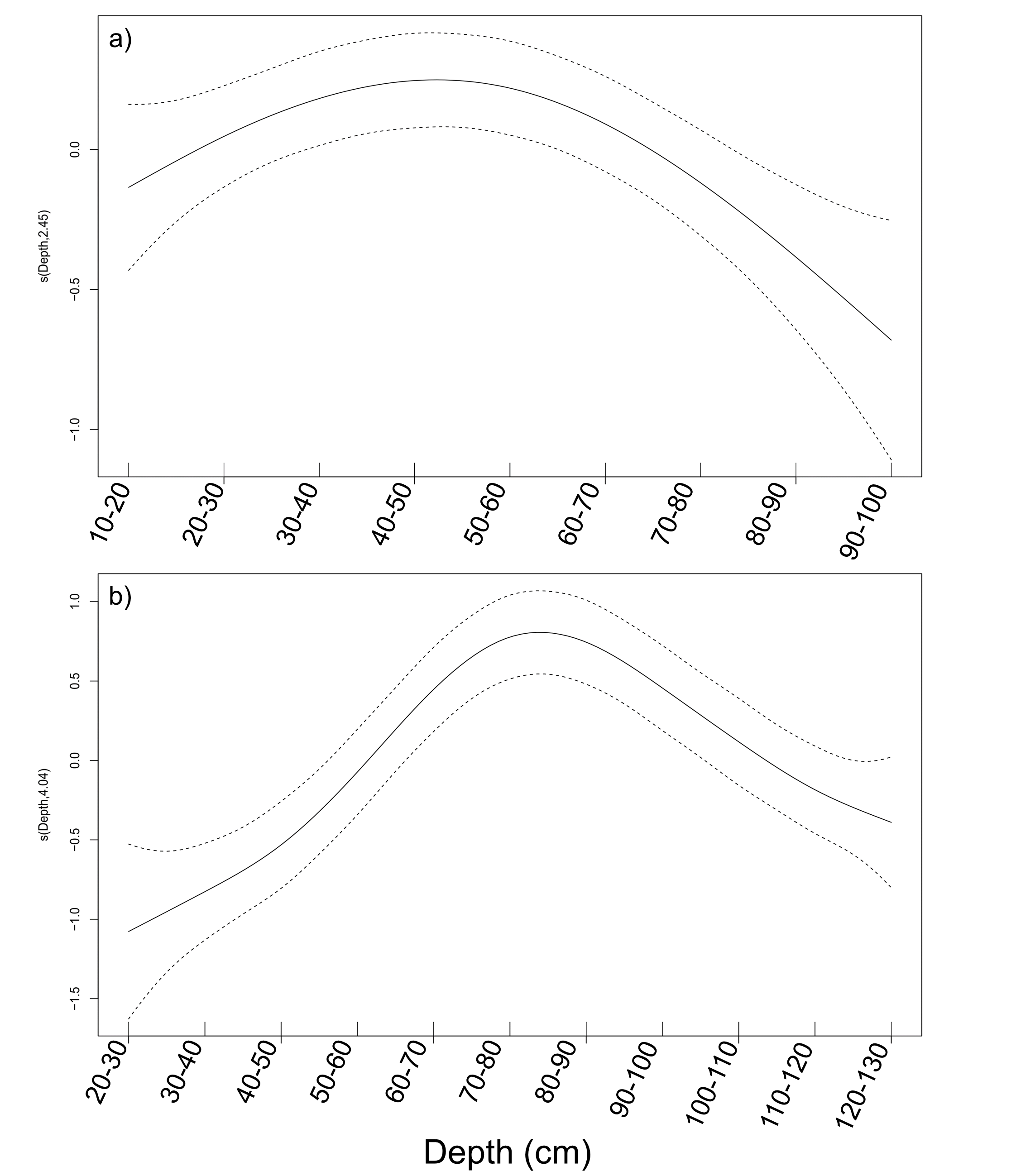

### Appendix S4

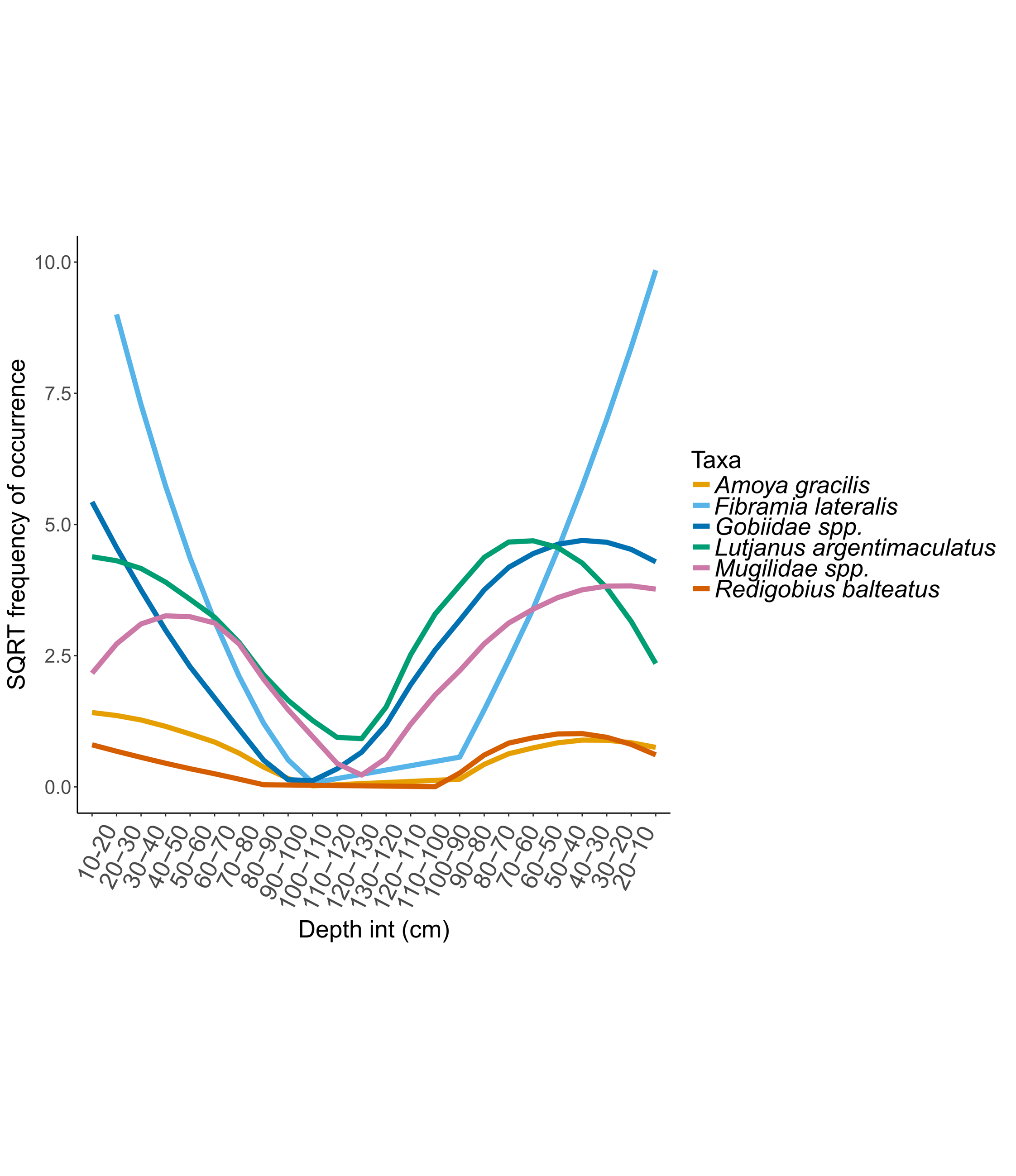

### Appendix S4

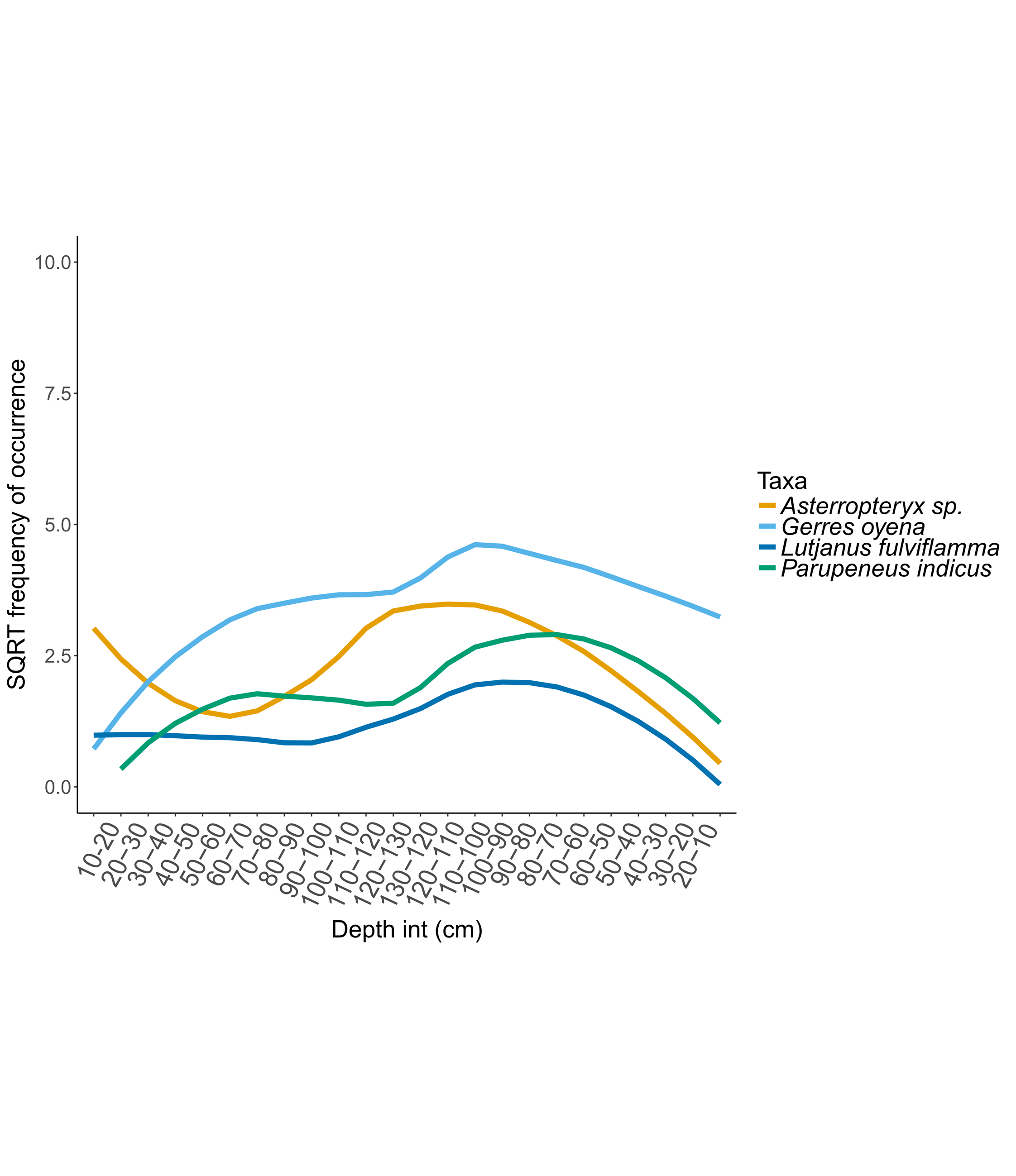

### Appendix S4

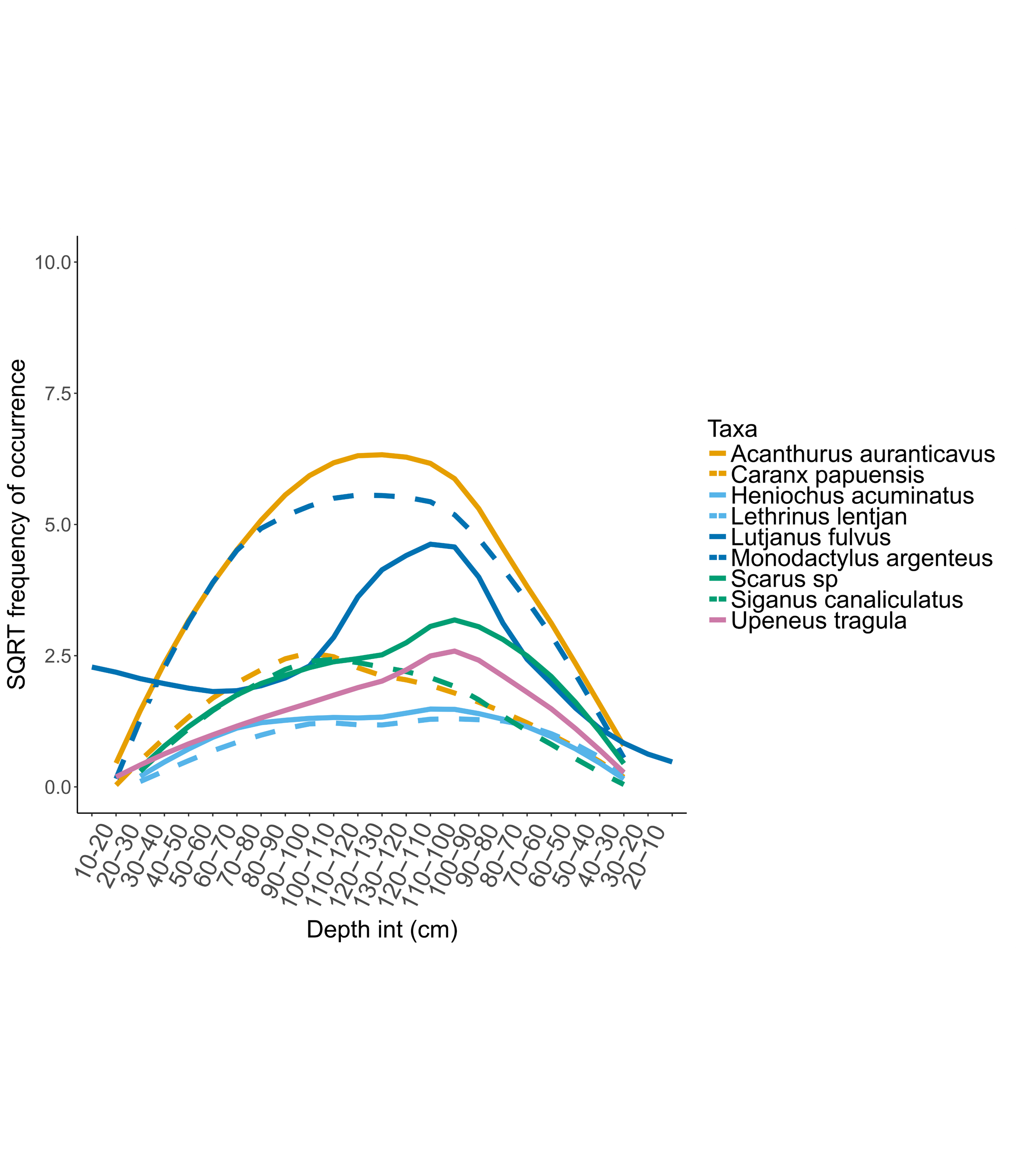

### Appendix S4

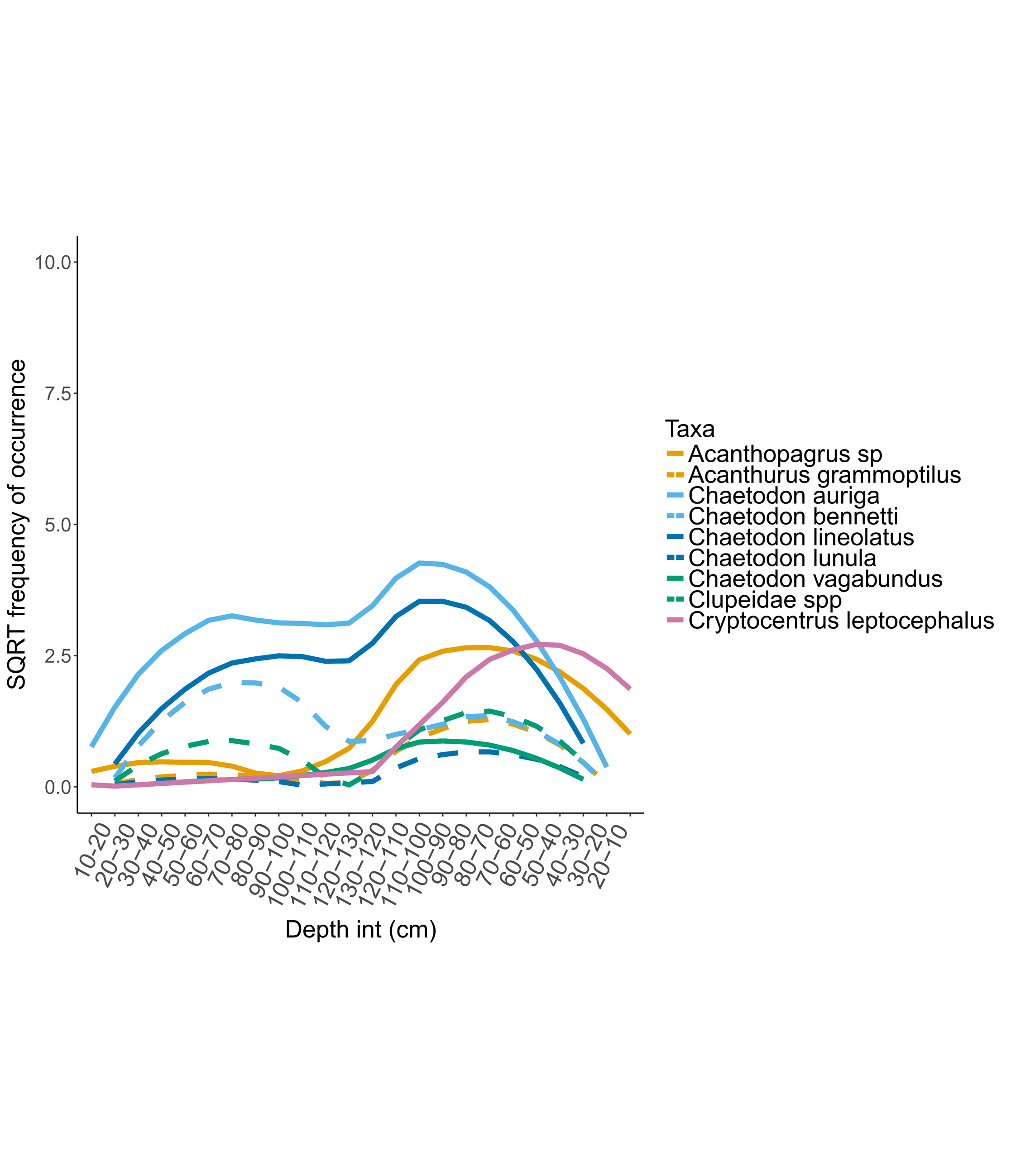

### Appendix S4

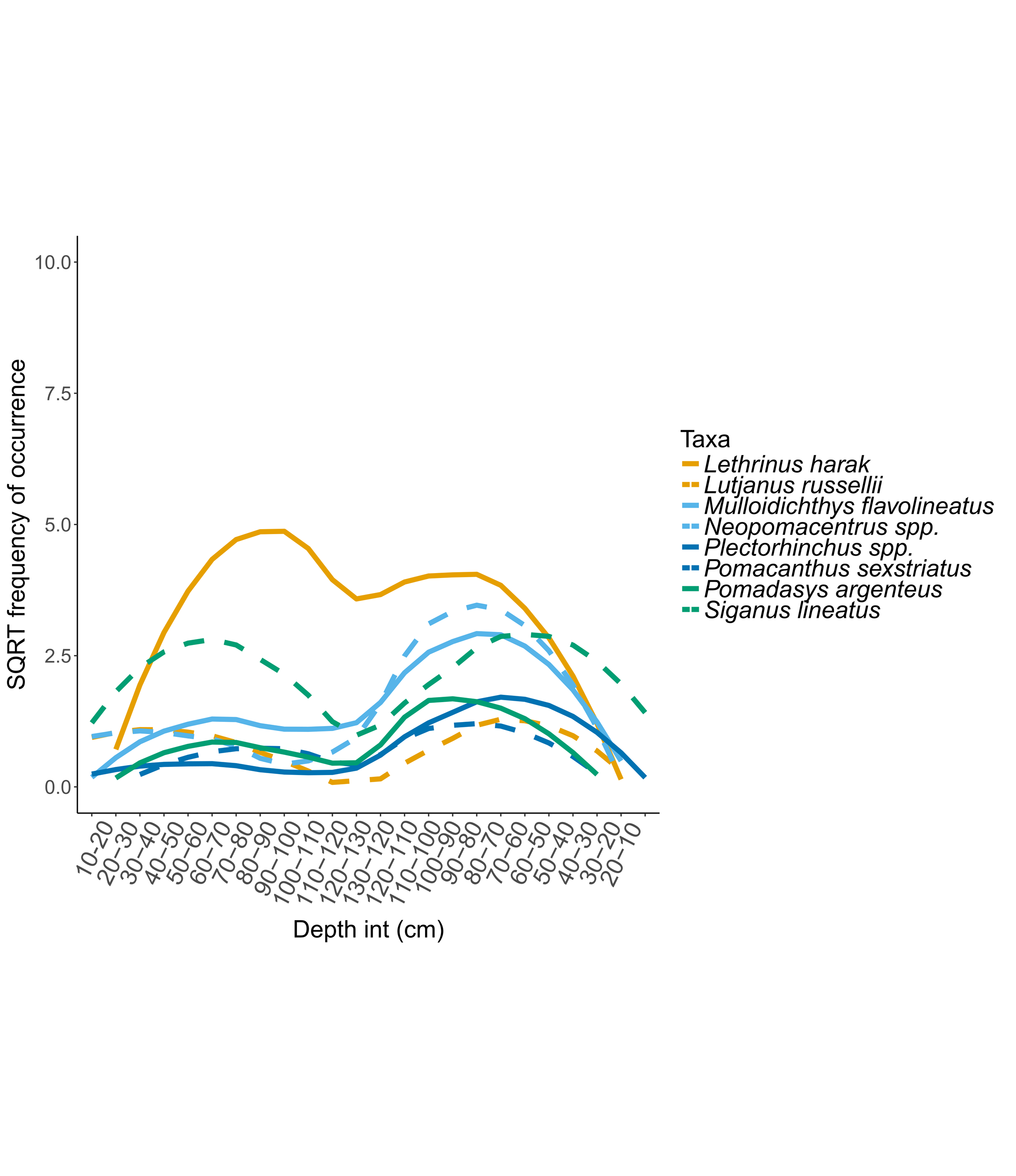
